# Uncoupling Neocortical Neuron Fate and Migration via a Let-7–RBX2 Axis

**DOI:** 10.1101/2025.09.11.675676

**Authors:** Steven Decker, Keiko Hino, Anna La Torre, Sergi Simó

**Affiliations:** Department of Cell Biology and Human Anatomy, University of California – Davis, Davis, CA, USA

**Keywords:** cortical development, neurogenesis, cell fate, let-7, RBX2

## Abstract

Throughout the central nervous system, the fate and migration of projection neurons are tightly coordinated to ensure that specific neuronal fates settle in precise spatial locations. This is particularly evident in the mammalian neocortex, where early-born projection neurons predominantly remain in the deeper layers of the cortical plate, whereas later-born neurons localize more superficially. However, it remains unclear whether neuronal fate acquisition directly primes the molecular mechanisms driving pyramidal neuron migration and positioning, or on the contrary fate and positioning are regulated independently. MicroRNAs have emerged as key regulators of cell fate determination in the neocortex. Among them, let-7 is known to influence neural progenitor competence and promote the neurogenesis of late-born projection neurons. Here, we show that let-7 also regulates projection neuron migration and positioning by targeting RBX2, a core component of the E3 ubiquitin ligase CRL5, which has been previously shown to inhibit neuron migration by terminating the Reelin/DAB1 signaling pathway. Let-7 directly binds to a conserved motif in the 3′ UTR of RBX2, reducing its translation and thereby diminishing CRL5 activity. Importantly, restoring RBX2 levels in the context of let-7 overexpression rescues the positioning of pyramidal neurons without altering let-7-induced effects on neuronal fate. Furthermore, we demonstrate that let-7 enhances pyramidal neuron migration by increasing locomotion speed and prolonging migratory activity. Together, these findings reveal that let-7 coordinates neuronal fate specification and migration via distinct molecular pathways, ensuring the proper laminar positioning of late-born pyramidal neurons in the neocortex.

## INTRODUCTION

The neocortex is a highly organized structure composed of six distinct layers of projection neurons (PNs) each characterized by specific connectivity patterns, molecular signatures, and functional roles ^1,2^. This intricate architecture emerges during development from the dorsal telencephalon, which initially is a thin pseudostratified epithelium comprised solely of neural stem cells. Over time, these stem cells differentiate into neural progenitors called Radial Glia Cells (RGCs) ^3,4^. Throughout corticogenesis, RGCs reside in the apical surface of the lateral ventricle (also known as ventricular zone (VZ)) and through a combination of direct and indirect neurogenesis, produce all PNs of the cortex ^5^.

As PNs are generated, their placement within the cortical layers follows a tightly regulated sequence. Newly born PNs migrate and settle on top of their predecessors in progressively more superficial layers, such that early-born PNs will reside in the lower cortical layers, whereas the later-born PNs will occupy in the upper layers ^3-5^. This spatial arrangement is not merely a byproduct of birth order but is tightly linked to neuronal identity: each PN subtype is specified to function within a particular cortical layer, and this functional specialization is inherently tied to its final position. This has been demonstrated by heterochronic transplant experiments showing that PNs can retain their laminar fate and position even when placed in a different temporal environment ^6,7^. Thus, there is a clear connection between the eventual fate of a PN and its migration dynamics to ensure that each cell is positioned in the correct layer. However, it remains elusive whether PN fate specification directly regulates PN migration and positioning or alternatively PN fate and position are independently regulated.

Recent studies suggest that both PN fate and migration are controlled, at least in part, by post-transcriptional mechanisms ^8-13^. Among these, microRNAs (miRNAs) have emerged as key regulators of both cell fate and migration during brain development ^14-18^. miRNAs are small non-coding RNA sequences that regulate gene expression by binding mRNA transcripts and promoting translational repression and mRNA degradation ^19^.

The generation of functional miRNAs involves a multistep biogenesis process. In the canonical biogenesis pathway, miRNAs are initially transcribed as long polycistronic transcripts and subsequently processed into precursor miRNAs (pre-miRNAs) by the Microprocessor complex. Pre-miRNAs are then exported into the cytoplasm, where the endoribonuclease Dicer cleaves them into mature, active miRNAs. Mature miRNAs associate with Argonaute (Ago) proteins to create the miRNA-induced silencing complex (miRISC), which recruits target mRNAs to promote translational repression and transcript degradation ^20^. Consequently, loss of Dicer disrupts the production of virtually all mature miRNAs ^19^. The critical role of miRNAs in brain development has been highlighted by the generation of conditional Dicer mutant mice ^21^. Dicer depletion disrupts cortical morphogenesis by affecting RGC competence and PN neurogenesis, and interfering with PN migration, ultimately leading to abnormal cortical layering and morphology ^14,15,22,23^.

Among the many miRNAs impacted by Dicer loss, the let-7 family has been identified as a key regulator of neural development. In the central nervous system, let-7 regulates cell fate decisions, cell cycle dynamics, and migration ^17,23-26^. In the cortex, let-7 expression has been detected in RGCs and PN, and increases on both cell types over embryonic development ^24^. Moreover, let-7 has been shown to drive changes in both cell fate and cell migration, and is sufficient to partially rescue the phenotype observed in Dicer conditional mutant animals ^23^. Mechanisms have been proposed for how let-7 regulates cell fate ^23^, but the mechanism(s) by which let-7 regulates cell migration are not understood.

The E3 ubiquitin ligase Cullin-5/RBX2 (CRL5) is a multiprotein complex involved in the regulation of neuronal migration in several structures of the nervous system ^27-30^. CRL5 is nucleated around the core proteins Cullin-5 (CUL5) and RBX2 (also known as RNF7) and recruits substrates for polyubiquitylation through different substrates adaptors proteins ^31^. During cortical development, CRL5 regulates PN migration by opposing to the Reelin/DAB1 signaling pathway ^27^. Reelin is an extracellular matrix protein secreted by Cajal-Retzius cells that triggers the phosphorylation of the intracellular adaptor protein DAB1 in PNs ^32^. In turn, DAB1 phosphorylation promotes PN migration by driving the switch from multipolar migration to locomotion at the boundary between the intermediate zone (IZ) and the cortical plate (CP), and triggering somal translocation at the top of the CP ^33-35^. As PNs reach the upper CP, expression of the CRL5 substrate adaptor SOCS7 increases, recruiting tyrosine-phosphorylated DAB1 (pY-DAB1) to CUL5/RBX2, and targeting it for degradation by the proteasome ^27^. CRL5-driven degradation of pY-Dab1 terminates Reelin/DAB1 signaling, ending PN migration. Depletion of Cul5, RBX2, or SOCS7, causes the accumulation of pY-DAB1, prolonged Reelin/DAB1 signaling, and PN over-migration into inappropriate cortical layers for their birthdate.

Here, we show that let-7 binds to a conserved motif in RBX2 3’UTR, reducing RBX2 protein expression and promoting the accumulation of DAB1, likely by decreasing CRL5 activity. Restoring RBX2 expression rescues let-7-induced change in PN positioning without altering fate specification, demonstrating that these two processes are molecularly separable. Finally, we used time-lapse microscopy to show that let-7 promotes PN migration by increasing migration velocity and sustaining migration for longer periods of time. Overall, our data indicate that let-7 is a master regulator that coordinates both the neurogenesis of late-born PN and their localization in the CP through different molecular mechanisms, and further demonstrate that PN fate and position can be independently regulated.

## Results

### Let-7 Targets RBX2 and Causes Dab1 Accumulation

To identify functionally relevant let-7 targets during PN migration, we first transfected a neural cell line (Ad12 HER10^36^) with either a let-7 mimic or a let-7 antagomir, a chemically modified oligonucleotide that specifically inhibits its target miRNA. RNA was collected two days after transfection and bulk RNA sequencing was performed (Figure S1A). Candidate let-7 targets were defined as transcripts that met the following criteria: 1) downregulated in cells treated with let-7 mimic and upregulated with let-7 antagomir, 2) predicted to contain conserved let-7 seed-matching sites in their 3′ untranslated region (3′ UTR) according to TargetScan ^37^, and 3) associated with gene ontology (GO) terms related to neuronal migration. This integrated approach yielded a single, high-confidence candidate: *RNF7* (RBX2, Figure S1B).

Conservation analyses indicated that let-7 binds to a highly-conserved sequence in the RBX2 3’UTR of several mammalian species (Figure 1A). To confirm that let-7 downregulates RBX2 mRNA levels, we transfected HER10 cells with a let-7 mimic or scrambled control. 48 hours post-transfection, RNA from transfected cells was collected and mRNA levels of RBX2 and two bona fide let-7 targets, HMGA2 ^23,38^ and Dicer ^18,39^, measured by RT-qPCR (Figure 1B). Upon let-7 expression, HMGA2, Dicer, and RBX2 mRNA levels decreased by 69.5±4.74%, 49.5±9.54%, and 57.6±8.5%, respectively. To determine whether let-7 directly targets RBX2 3’UTR, we performed a dual luciferase reporter assay. The full RBX2 3’UTR sequence was cloned downstream of a firefly luciferase coding sequence in an expression plasmid. A plasmid expressing *Renilla* luciferase was co-transfected in all our experiments and used as an internal control for normalization. The resulting firefly luciferase-RBX2 3’UTR plasmid (namely RBX2 3’UTR) was co-transfected in tsa201A cells with or without a let-7a expression plasmid. Co-expression of let-7 led to a 27.2% decrease in firefly luciferase compared to the control (Figure 1C). No change in signal was observed when a different miRNA (miR-7) was co-expressed or when a firefly luciferase plasmid without the RBX2 3’UTR sequence (Empty 3’UTR) was co-transfected with let-7. These data indicate that let-7 recruits RBX2 3’UTR and targets the mRNA for degradation. Next, we tested whether let-7 affects RBX2-dependent CRL5 activity. To test this end, we transduced primary mouse cortical neurons with lentiviral particles overexpressing let-7 and observed a decrease in RBX2 levels by western blotting when let-7 was expressed (Figure 1D and 1E). Importantly, decreased RBX2 levels were accompanied by an accumulation of DAB1, suggesting that let-7 decreases CRL5 activity by opposing RBX2 expression. Collectively, these data suggest that let-7 directly interacts with the 3’UTR of RBX2, opposes RBX2 protein levels, and reduces CRL5 activity in PNs.

**Figure 1:**
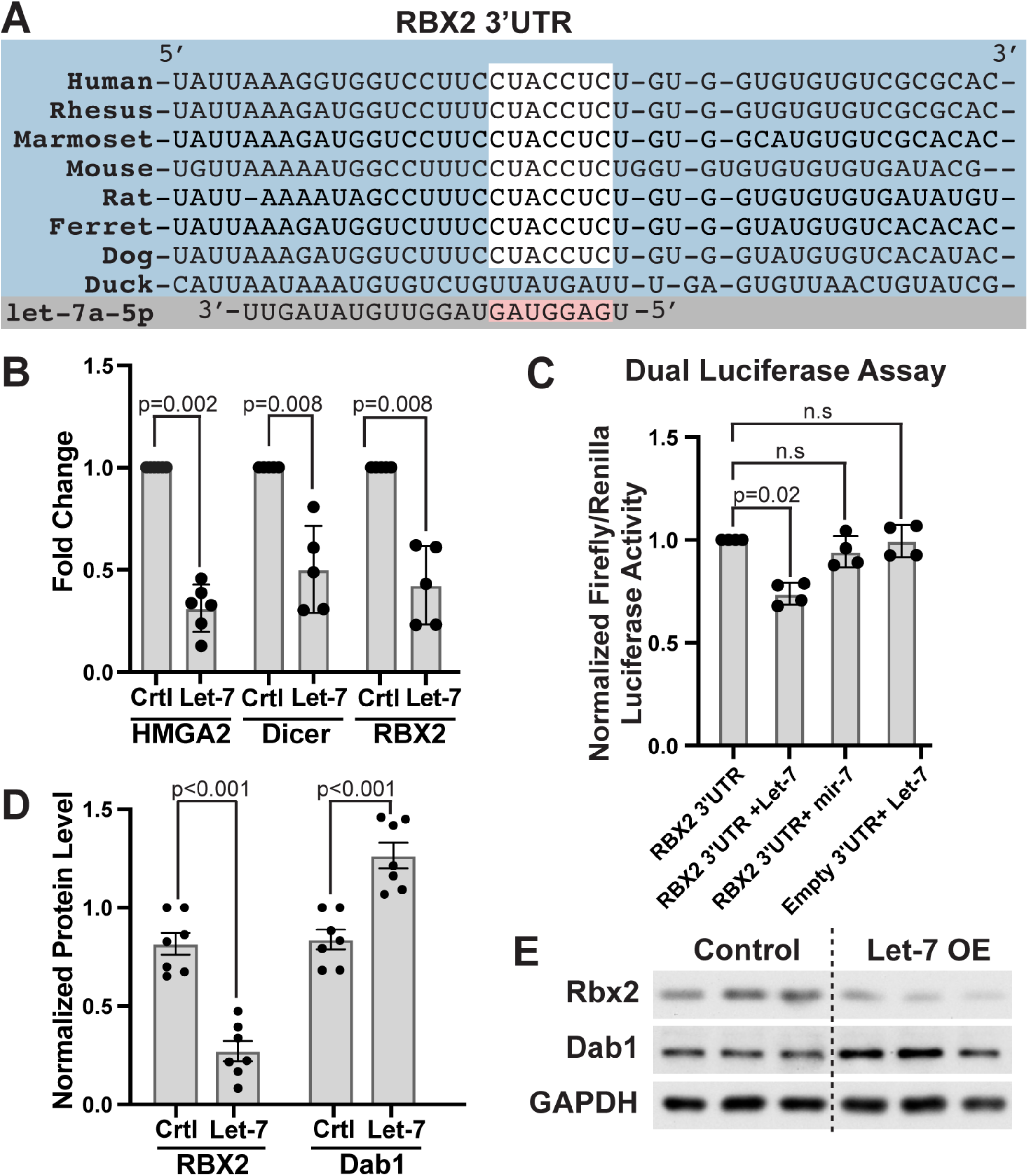
Let-7 directly binds RBX2 3’UTR and downregulates CRL5 activity. (A) Diagram highlighting let-7a-5p seed sequence (pink) and its target sequence conserved in multiple mammalian RBX2 3’UTR sequences (white box). (B) RT-qPCR fold change of indicated genes in HER10 cells transfected with a let-7 mimic or a scrambled control (Ctrl). Mean ± SD. Statistics, Mann-Whitney test. (C) Dual luciferase assay in HEK293 cells transfected with a let-7 mimic, a negative control (mir-7), or empty 3’ UTR is equivalent plasmid with no UTR insert. Mean ± SD. Statistics, Kruskal-Wallis test with Dunn’s multiple comparison test. (D-E) Western blot of primary mouse cortical neurons transduced with lentiviral particles over-expressing let-7 (Let-7 OE) or scramble control (Control) and quantification of RBX2 and DAB1 Western blots. Mean ± SD. Statistics, Welch’s t test.

### RBX2 Rescues Let-7 Induced Overmigration

Previous work from our group and others has shown that overexpression of let-7 causes PN to localize more superficially in the cortical plate (CP) than expected based on their time of birth^23,29^, indicating that let-7 promotes PN overmigration. We next asked whether this phenotype results from let-7-dependent repression of RBX2 levels and subsequent inhibition of CRL5 activity. To test this hypothesis, we overexpressed let-7 at embryonic stage (E) 13 by *in utero* electroporation (IUE) together with a fluorescent mCherry reporter, and we examined the position of the electroporated cells at postnatal day (P) 0. An empty plasmid was used as control electroporation. To quantify the position of the electroporated cells, the CP was divided into eight bins and the percent of electroporated PNs (mCherry fluorescent positive cells) in each bin was quantified using the RapID automatic counter ^40^. As expected, upon let-7 overexpression, we observed an overmigration of the electroporated cells with more cells positioned in the upper cortical layers (bins 1 and 2) (Figure 2A, B, and D). Next, we co-electroporated let-7 with a plasmid expressing the cDNA of RBX2, which lacks the 3’UTR, and therefore the let-7 target site. Strikingly, RBX2 co-expression resulted in a full rescue of PN overmigration when compared to let-7 expression alone (Figure 2B-D). To rule out that RBX2-dependent changes in PN position were a consequence of changes in PN fate, we quantified the percent of electroporated cells that are Brn2+/Ctip2-, which identifies late-born PNs ^2^. As expected, let-7 expression resulted in a significant increase in the proportion of Brn2+/Ctip2-PNs. Co-expressing RBX2 did not cause any fate changes in comparison to let-7 expression alone (Figure 2E). Moreover, there was no change in PN position or fate when RBX2 cDNA was electroporated alone (Figure S2A-C). We also tested whether RBX2 depletion regulates PN fate. To that goal, we generated conditional RBX2 knockout animals (RBX2cKO) by crossing Emx1-Cre mice with a RBX2^flox/flox^ line, resulting in loss of RBX2 expression in the dorsal telencephalon beginning around embryonic day (E) 10, as reported before ^29^. These animals exhibit significant layering defects but had normal proportions of CUX1+, CTIP2+, and FOXP2+ PNs (Figure S2D-E). Collectively, these data suggest that let-7 regulates PN localization by targeting RBX2 and this mechanism is independent of the let-7 role in PN fate determination.

**Figure 2:**
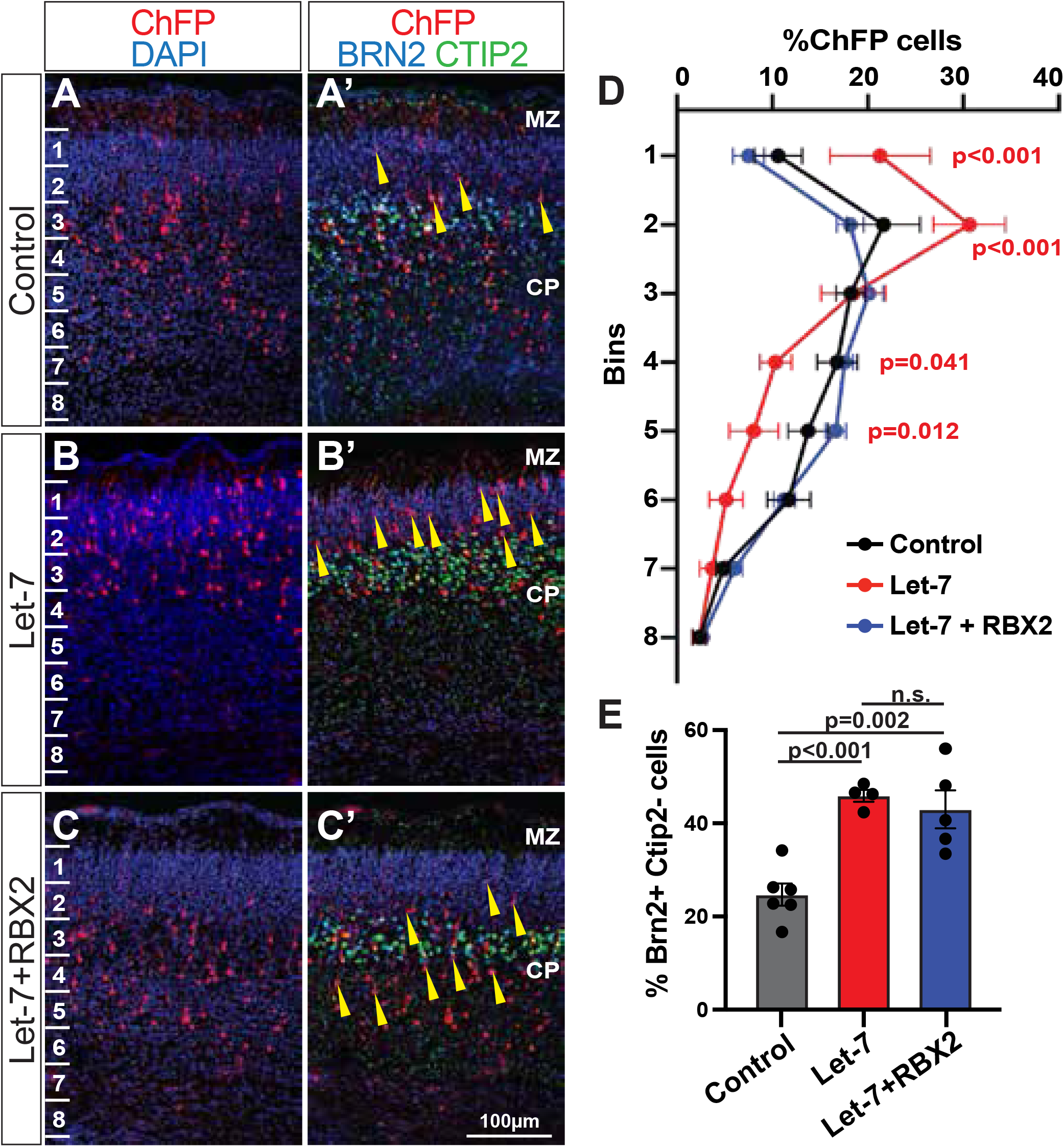
RBX2 expression rescues migration but not fate defects caused by let-7 overexpression. (A-C’) Representative images of cortices electroporated at E13 and collected at P0 showing PNs (ChFP+) expressing control (n=10), let-7 (n=5), or let-7+RBX2 (n=7) plasmids. Sections were counterstained with DAPI (A,B, C) and stained for BRN2 (blue) and CTIP2 (green) (A’, B’, C’). Yellow arrowheads indicate BRN2-positive, CTIP2-negative, electroporated (ChFP) PNs. (D) Percentage of control, let-7, and let-7+RBX2 electroporated PNs (ChFP+) in each indicated bin. Mean ± SD. Statistics, p values represent comparison between control (A-A’) and let-7 (B-B’) electroporations using two-way Anova with Tukey’s multiple comparison test. Let-7+RBX2 (C-C’) was not significantly different from control (A-A’) in any bin. (E) Percentage of electroporated cells from control (A-A’), let-7 (B-B’), and let-7+RBX2 (C-C’) that showed positively stained for BRN2 (BRN2+) and showed no staining for CTIP2 (Ctip2-) after immunofluorescence. BRN2+/CTIP2-were considered upper-layer PNs. Mean ± SD. Statistics, one-way Anova with Tukey’s multiple comparison test.

### Let-7 regulates the speed and path of migrating PNs

To understand how let-7 is regulating final PN position in the CP, first, we analyzed the migration behavior of PNs expressing let-7. Let-7 was overexpressed by IUE at E13.5 as described above, and brains we collected at E15.5 to capture actively migrating PNs, enabling direct observation of let-7’s effect during a critical developmental window. Electroporated brains were sliced using a vibratome and fluorescent PNs in organotypic brain slices were imaged for 18-24 hours (Figure 3A and Movie S1). Interestingly, the PNs overexpressing let-7 had a modest, yet consistent higher average migration speed (15.7±5.7 µm/hr, N=62 cells) compared to control cells (12.7±3 µm/hr, N=49 cells, Figure 3B). To measure the path linearity of the migrating cells, we measured the ratio of their displacement to the total distance traveled. Control PNs had a linearity ratio of 0.69 while the let-7-overexpressing PNs had a ratio of 0.75 (Figure 3C), indicating that let-7 enhances both the efficiency and directionality of PN migration. Next, we assessed whether the differences observed in PN migration could be attributed to changes in the morphology of the RGCs surrounding the electroporation site. Control and let-7 expressing IUEd brains were stained against the RGC marker Nestin ^41^. We observed no change in morphology of RGCs between control and let-7 electroporations (Figure S4), supporting the idea that let-7 acts in a cell-autonomous manner to regulate PN migration. Collectively, these data show that let-7 fine-tunes PN migration by modulating both the speed and trajectory of cells during locomotion.

**Figure 3:**
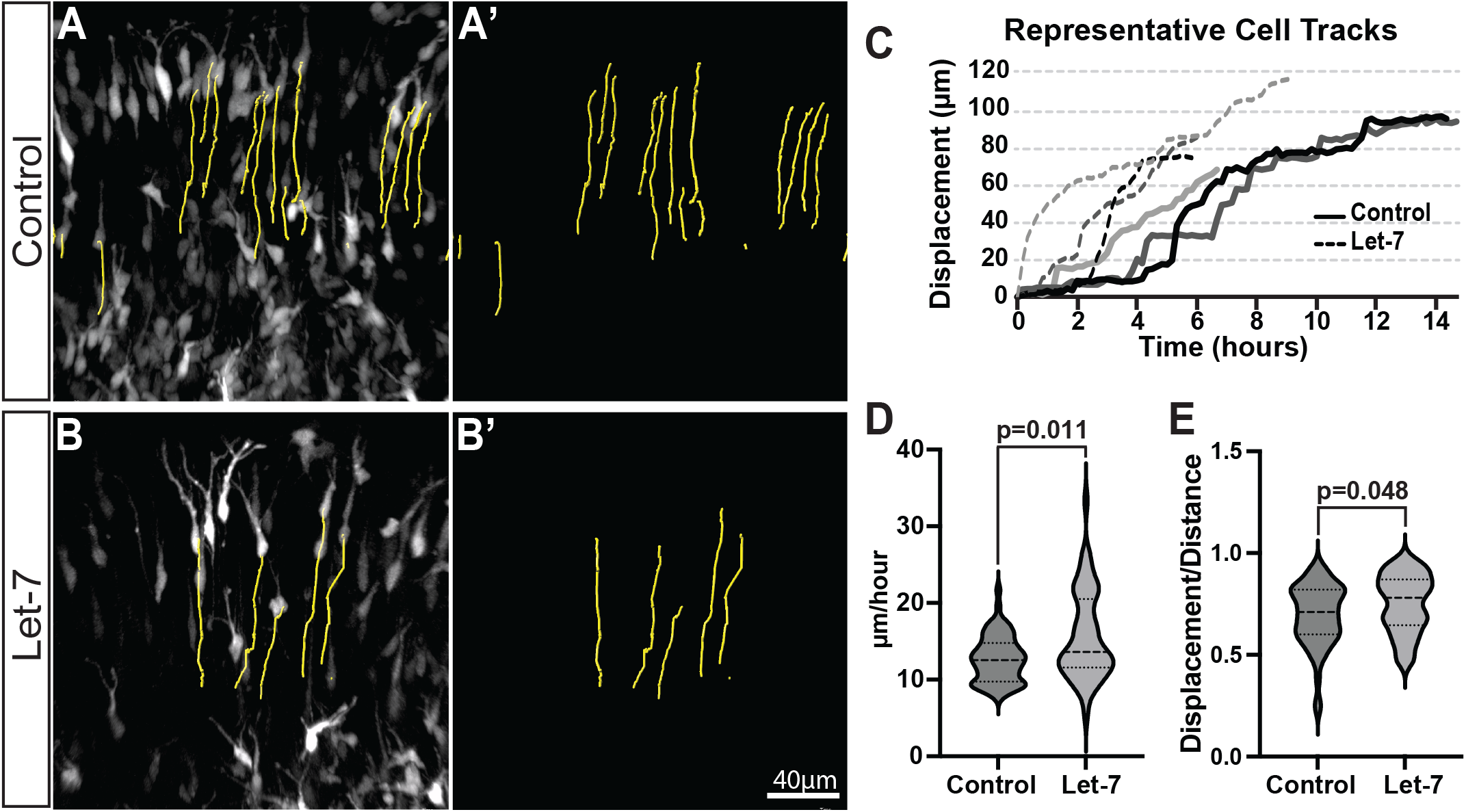
Let-7 regulates cortical pyramidal neuron migratory behavior. (A-B’) Portion of still images from supplemental movies showing cortices electroporated at E13 with control (A-A’) or let-7 (B-B’) expressing plasmid and collected for time-lapse imaging at E15. Yellow lines represent cell tracks of individual PNs. (C) Tracking of three control (solid lines) and let-7-expressing (dashed lines) PNs over time. Line length represent the time that cell was tracked. (D-E) Violin plots representing control (n=62) and let-7 (n=49)-expressing PNs migration speed (D) determined by averaging the distance completed by a single PN in all dimensions during the time that the cell was tracked and cell track straightness (E) by measuring the ratio of displacement to track length. Violin plot data: dashed line, median; dotted lines, 25th to 75th percentile. Statistics, Mann-Whitney test.

### Let-7 Regulates Duration of Migration

To understand the long-term migration behavior of let-7-expressing PNs, we performed IUE using control or let-7 expressing plasmids at E13. Electroporated brains were collected daily from E14 until birth. We confirmed that let-7 over-expression did not alter RGC morphology, ensuring that potential migration alterations were not a secondary effect of a defective RGC (Figure S3). At E14, all the electroporated cells were in the VZ/SVZ in the control and let-7 conditions indicating that let-7 does not alter early migration steps (Figure 4A-C). Between E15 and E17, the electroporated cells in both conditions have entered the CP and the proportion of PNs present in the VZ/SVZ/IZ and CP is similar (Figure 4D-G and 4H-K). By E17, most control PNs have reached the top of the CP and ended migration. In contrast, let-7-expressing PN sustained migration, remaining closer to the top of the CP in comparison to control PNs (Figure 4L-O). This overmigration phenotype was also observed at E18 and P0 (Figure 4P-S and Figure 2, respectively). Notably, the overall proportions of PNs within the CP versus IZ/SVZ/VZ remained consistent between groups at all stages examined, indicating that let-7 does not alter the general distribution but specifically extends the duration of PN migration.

**Figure 4:**
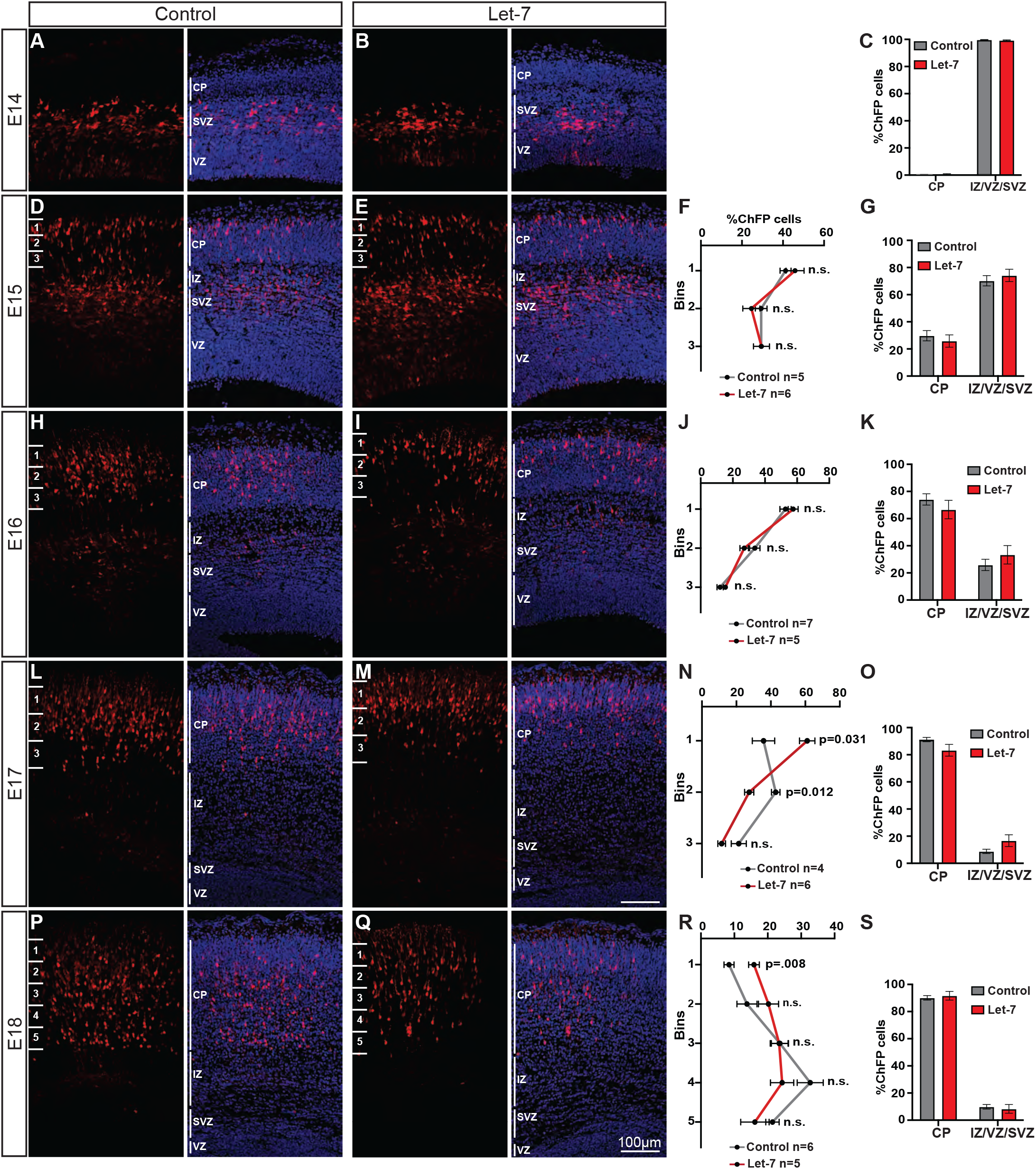
Let-7 prolongs neuronal migration. Control or let-7-expressing plasmids were in utero electroporated into E13 embryos and collected at E14 (A-B), E15 (D-E), E16 (H-I), E17 (L-M), and E18 (P-Q). Brain slices were counterstained with DAPI. No differences were observed in the percentage of electroporated neurons (red) within the CP and the IZ/SVZ/VZ regions between let-7-expressing and control neurons in the different ages analyzed (C, G, K, O, S). Mean ± SD. Statistics, Welch’s t test. Distribution of electroporated neurons within the CP showed that let-7 expression did not affect neuron distribution at earlier stages (F, J), but sustained the migration of neurons towards more upper layers (N, R). Mean ± SEM. Statistics, Student t test with Bonferroni-Dunn’s multiple comparison test.

## Discussion

The sequential nature of cortical PN generation intimately associates birthdate to both laminar identity and final laminar position ^42^. Since the landmark heterochronic transplant experiments pioneered by Dr. Susan McConnell, evidence has supported that PN birthdate strongly predicts laminar position, even when cells were placed in environments with differently aged neurons. These findings also indicate that fate and positioning are largely governed by cell-autonomous mechanisms albeit environmental factors might also participate in fine tuning PN position ^6,7^.

Despite this, it remains unclear whether PN fate determination directly regulates laminar position or whether fate and position are regulated by parallel but independent mechanisms. Supporting the latter hypothesis, we propose a model whereby let-7 acts as a master regulator of late-born PNs by coordinating both their fate specification and location in upper cortical layers during development. Let-7 expression increases throughout development of the cortex, both in RGCs and PNs ^24^. Thus, elevated levels of let-7 in RGCs contribute to the neurogenesis of late-born PNs, even at times when earlier-born PNs are also being generated ^23^. Us and others have previously showed that ectopic expression of let-7 in RGCs during early or mid-development stages drives a precocious neurogenesis of late-born PNs (Figure 2)^17,28^. Importantly, these “early” late-born PNs localize in the upper layers along wild type PNs born at later stages. But how do late-born PNs, whether endogenous produced or ectopically generated, reach their appropriate upper cortical layers? We propose that higher let-7 levels downregulate the expression of RBX2, thereby decreasing CRL5 activity. This downregulation accelerates and sustains PN migration, promoting the correct laminar positioning of late-born cells.

Let-7 regulates PN fate at least in part by downregulating the high-mobility group AT-hook 2 (HMGA2) ^23^. HMGA2, a chromatin-associated factor, is known to promote stem cell self-renewal and proliferation, but no role on PN migration has been described. As development progresses, the increase of let-7 downregulates HMGA2 and promotes neurogenic competence and differentiation of RGCs.

Extending beyond its role in fate specification, our findings reveal that let-7 also modulates PN migration through a distinct mechanism: by directly targeting RBX2, leading to reduced CRL5 activity, accumulation of DAB1, and enhanced migration speed and persistence ^27,43,44^. This aligns with previous reports showing that CRL5 regulates the speed and duration of migration by opposing Reelin/DAB1 signaling ^27,44^. Let-7 also regulates the linearity of the PN migration path, suggesting broader effects on migration dynamics. Importantly, these effects do not seem linked to changes in the surrounding RGC scaffold, pointing to cell-autonomous mechanisms. Our sequencing data also identifies seven other let-7 target genes, besides RBX2, listed in the gene ontology term “Regulation of Cell Migration” (GO: #0030334) (DUSP22, SLK, KIF2A, INSR, TGFBR1, IGFR1, and ROBO1). Although we cannot exclude the possibility that the observed let-7 effects on PN migration could be a consequence of altering any of these other genes, we reason that our experiment rescuing let-7-induced PN overmigration by solely expressing RBX2 suggest that let-7 regulation of CRL5 activity is mainly responsible for the phenotypes observed. Alternatively, it is possible that the migration effects may be due to changes in let-7-dependent, CRL5-independent, PN polarity or their adhesion to the RGCs ^45,46^ and future work will address these possibilities.

MiRNAs are well-established regulators of neurogenesis and different aspects of neuronal migration ^47-50^. For example, miR-9 targets both fate determinants, such as HES1 ^51^, FOXG1 ^52^, and FOXP2 ^53^ and cytoskeletal regulators, including MAP1B ^54^ and N-Cadherin ^55^. Similarly, miR-124 influences neural differentiation through the REST complex ^56^ and has also been implicated in modulating small GTPases and other cytoskeletal regulators ^57^. miR-128, in turn, regulates the Reelin pathway ^58^ and neuronal migration more broadly ^49^, though it is primarily expressed during early corticogenesis and downregulated at later stages ^23^. Notably, however, let-7 remains unique in that it has been shown to regulate both cell fate and migration within the same population, highlighting its multifaceted role in cortical development.

Overall, our study identifies let-7 as a key coordinator of cortical development that independently regulates neuronal fate and migration by targeting distinct molecules, HMGA2 for fate ^23^ and RBX2 for migration (this paper). By modulating CRL5 activity through RBX2 repression, let-7 fine-tunes migration dynamics critical for proper cortical layer formation. Disruptions in this regulatory axis may underlie neurodevelopmental disorders characterized by impaired neuronal migration and cortical organization, such as epilepsy and intellectual disability. Moreover, altered let-7 expression has been implicated in neuropsychiatric conditions including autism spectrum disorders and schizophrenia ^59,60^, where disrupted cortical layering and connectivity are common features. Our work therefore not only advances fundamental understanding of cortical development but also provides a molecular framework with potential implications for diagnosing and therapeutically targeting neuronal migration disorders.

## Supplemental Figure Lengends

**Figure S1: Let-7 targets in HER10 cells.**

(A) RNA-seq analysis comparing HER10 cells transfected with a let-7 mimic (n=5) or a let-7 antagomir (n=5).

Ratio of counts per million between let-7 mimic and let-7 antagomir is plotted. The x axis represents the logarithmic fold ratio of let-7 antagomir/let-7 mimic per gene identified. The y axis represents the negative logarithmic (-log_10_) adjusted p value (-log Adj. P val) calculated by using the Benjamini-Hochberg false discovery rate method. Only genes with an average count per million (cpm) of >0.5 were used for statistical purposes and plotted. Purple dots represent genes that are predicted let-7 targets using TargetScan.

(B) Gene ontology term classification of significant differentially expressed genes between between let-7 mimic and let-7 antagomir HER10 samples. In the “Regulation of Neuron Migration” (GO:2001222) only one gene was identified: RBX2.

**Figure S2: RBX2 does not participate in cell fate specification in the neocortex.**

(A) Representative immunofluorescence images for BRN2 and CTIP2 in P0 brain slices from embryos in utero electroporated at E13. Embryos were electroporated with control or RBX2-expressing plasmids. Yellow arrowheads indicate BRN2 positive, CTIP2 negative, electroporated (red) cells.

(B) Percentage of BRN2 positive, CTIP2 negative, in control and RBX2-expressing neurons. Mean ± SD. Statistics, Welch’s t test; n.s., not significant.

(C) Distribution of control (n=3) and RBX2 (n=3)-expressing electroporated in the cortical plate at P0. No changes in cell position were observed between conditions. Mean ± SEM. Statistics, Student t test with Bonferroni-Dunn’s multiple comparison test.

(D) Representative images of control (RBX2^fl/fl^) and Rbx2 conditional mutant (RBX2^fl/fl^; EMX1-CRE) cortices at P0 stained against CUX1, CTIP2, and FOXP2.

(E) Quantification of Cux1, CTIP2, or FOXP2 number of pyramidal neurons in control (RBX2^fl/fl^) and Rbx2 conditional mutant (RBX2cKO). No changes in cell fate specification were observed. Mean ± SEM. Statistics, Student t test with Bonferroni-Dunn’s multiple comparison test; n.s., not significant.

**Figure S3: Let-7 overexpression does not disrupt RGC morphology.**

Representative immunofluorescence images for Nestin at E14 (A-B’) and E15 (C-D’) in brain slices from embryos electroporated in utero at E13 with either control (A–A’, C–C’) or let-7 (B–B’, D–D’)-expressing plasmids.

## Materials and Methods

### Animals

All animals were used with approval from the University of California Davis Institutional Animal Care and Use Committees and housed and cared for in accordance with the guidelines provided by the National Institutes of Health. RBX2 floxed mice (RBX2^fl/fl^) and Rbx2^fl/fl^:Emx1-Cre (RBX2cKO) were generated as previously described ^27,29^.

### Cell Culture

#### Primary cultures

For primary cultures, E16 cortices were dissected and pooled from a single litter and dissociated using a Papain dissociation system (Worthington, LK003150) following the manufactures instructions. 4 million cells were plated into 35mm wells pre-coated with 0.05mg/ml Poly-D-Lysine (Sigma-Aldrich, P6407). Cells were maintained in Neurobasal medium (Life technologies, 21103049), with Glutamax (Life Technologies, 35050061), B27 supplement (Life Technologies, 17504044), Penicillin/Streptomycin (Life Technologies, 16140122), and glucose at a final concentration of 0.55%.

#### Viruses

Lentiviruses were generated with the help of the National Eye Institute Core Facility - UC Davis Viral Core using pLenti-GFP (control) and Lv-let-7d-5p (let-7). LV-Let7d-5p was a gift from Michele De Palma (Addgene plasmid # 117055 ; http://n2t.net/addgene:117055 ; RRID:Addgene_117055) ^61^. 293T cells were co-transfected with the 2^nd^ generation of lentiviral plasmids, psPAX2, pMD2.G, and control or let-7 plasmids. 72 hours later, the media was collected, viruses were concentrated by ultracentrifugation, and stored at −80°C. Primary neurons were transduced 48hr after plating using a mixture of 800µl complete medium and 30µl of viral particles for 5 hours. After 5 hours, media was changed, and cells were collected 72 hours later.

#### RNA sequencing

HER10 cells were transfected with a Let-7 mimic or antagomiR (Dharmacon C-300478-07-0010 and IH-3004478-08-0010) at a final concentration of 10nM using Lipofectamine 2000 (Fisher Scientific, 11668019) following the manufactures instructions. Two days later cells were collected and total RNA from let-7 mimic (n=5) and let-7 antagomiR (n=5) cells was extracted using the Total RNA Purification Plus Kit (Norgen Biotek Corp.,48300). Gene expression profiling was carried out using a 3’-Tag-RNA-Seq protocol. Barcoded sequencing libraries were prepared using the QuantSeq FWD kit (Lexogen) for multiplexed sequencing according to the recommendations of the manufacturer using both the UDI-adapter and UMI Second-Strand Synthesis modules (Lexogen). The fragment size distribution of the libraries was verified via micro-capillary gel electrophoresis on a LabChip GX system (PerkinElmer). Libraries were quantified by fluorometry on a Qubit instrument (Life Technologies), and then pooled in equimolar ratios. The library pools were quantified by qPCR with a Kapa Library Quant kit (Kapa Biosystems/Roche). Finally, the library pool was sequenced on a HiSeq 4000 sequencer (Illumina) with single-end 100 bp reads. Raw data and differential gene expression analyses are available at Gene Expression Omnibus (GEO) repository (NIH) by the time of publication.

#### Luciferase Assay

Luciferase assays were performed using GeneCopoeia Luc-Pair Duo-luciferase Assay Kit 2.0 (GeneCopoeia LF001). HEK293 cells were co-transfected with mouse RNF7 (RBX2) 3’UTR target expression clone or control vector with no 3’UTR insert (GeneCopoeia MmiT094746-MT06 and CmiT000001-MT06) and let-7 mimic or control mimic (miR-7) at a final concentration of 10nM. 24hrs later the cells were lysed and luciferase activity was measured following the manufactures instructions.

#### qPCR

HEK293 cells were transfected with let-7 mimic at a final concentration of 10nM. RNA was collected 48 hours later using TRIzol reagent (Life Technologies, 15596018) and cDNA was generated using iScript cDNA synthesis kit (Biorad, 1708890). qPCR was performed using SYBR green power PCR mastermix (life technologies, 4367659) and an Applied Biosciences StepOne qPCR system. Transcript abundance was quantified using the ΔΔCt method. For primer sequences find Primer sequence table.

#### In *Utero* Electroporation

In *utero* electroporation was performed at E13.5 as previously described ^27,29,44^ using timed pregnant CD-1 mice. DNA solutions containing 0.5µg/µl pCAG-EGFP (control) or 0.5µg/µl pCAG-mCherry and 1µg/µl Lv-let-7d-5p (Let-7 OE) were mixed in 10mM Tris pH 8.0 and 0.01% Fast Green. For rescue experiments, 0.5µg/µl pCAG-mCherry, 1µg/µl Lv-let-7d-5p (Let-7 OE), and 0.125µg/µl pCAG-T7-RBX2 in the same injection solution was used. A 0.5µg/µl pCAG-mCherry and 0.125µg/µl pCAG-T7-RBX2 mix was used to express RBX2 alone. pCAG-T7-RBX2 expresses a T7-tagged mouse RBX2 cDNA, obtained from embryonic brain mRNA by a reverse transcription reaction (SuperScript II Reverse Transcriptase system, Invitrogen) using Oligo-dT (Invitrogen) primer and PCR amplification (Herculase II, Agilent Technologies). Of note, pCAG-T7-RBX2 expresses a heterologous 3’UTR with no let-7 target sequence. 1 µl of the solution was microinjected into the lateral ventricle of each embryo. Tweezerodes electrodes (BTX) with 5mm pads were used for electroporation with five 50ms-pulses of 30V. Electroporated brains were collected between E14.5 and P0 as specified in the text.

#### Live Imaging

In *utero* electroporation was performed with let-7 and control conditions on E13.5 embryos as described above. Electroporated embryos were dissected two days later in cold media containing: 248mM sucrose, 1mM CaCl_2_, 5mM MgCl_2_, 10mM glucose, 4mM KCl, 26mM NaHCO_3_, and 0.5% phenol red. Brains were imbedded in 3% low melting point agarose (VWR International, 89125-532) in Neurobasal media (Life technologies, 12348017). 300µm coronal sections were cut using a Leica VT1000 vibratome and sections were cultured on a transwell (Fisher scientific, PICM03050) in a glass-bottom culture dish (World Precision Instruments, FD5040) with DMEM/F12 media (Life technologies, 11039021) supplemented with 10% FBS, Penicillin/Streptomycin, and B27. Organotypic cultures were cultured at 37°C with 95% O_2_ + 5% CO_2_ in a Dragonfly spinning disc confocal microscope outfitted with an incubator. Z-Stacks were taken every 10 minutes for 18-24 hours using a Leica HC FLUOTAR L 25x water-immersion objective. Tissue drift was corrected, and cell tracks were quantified using Imaris (Oxford Instruments).

#### Immunofluorescence

After dissection, brains were immediately placed in 4% Formalin (Fisher Scientific, 23245684) at 4°C overnight. The next day, samples were cryoprotected with a 30% sucrose/PBS solution and kept at 4°C overnight. Brains were frozen in a 1:1 mixture of 30% sucrose/PBS and OCT compound (Fisher Scientific, 23730571) and 14µm sections were cut using a cryostat. Sections were blocked with 10% normal donkey serum in PBS with 0.3% Triton-X for 1hr at room temperature. Primary antibody (see antibody table) incubations were done in blocking buffer at 4°C overnight and secondary antibody incubations were done with blocking solution at room temperature for 1hr. Sections were counterstained with DAPI and mounted using Flouromount-G mounting media (Invitrogen, 00-4958-02). Slides were imaged using an Olympus FV3000 or FV4000 laser scanning confocal microscope.

#### Western blotting

For western blotting samples were lysed with lysis buffer (50 mM Hepes, pH7.5, 150 mM NaCl, 1.5 mM MgCl2, 1 mM ethylene glycol tetraacetic acid, 10% glycerol, and 1% Triton X-100). Protease inhibitors (Sigma-Aldrich, P1860) and phosphatase inhibitors(10 mM NaF, and 1 mM Na3VO4) were added to the lysis buffer just prior to homogenization. After lysis, samples were centrifuged at 15,000g for 15 minutes at 4°C. Cleared supernatant was denatured at 98 °C for 10 min in sample buffer (10 mM Tris, pH 6.5, 150 mM β-mercaptoethanol, 0.5% sodium dodecyl sulfate (SDS), 5% glycerol, and 0.0125% bromophenol blue). Samples were run on a tris-glycine gel and transferred to a nitrocellulose membrane (BioRad, 1620112). Membranes were blocked in 5% nonfat milk in TBST for 1 hour at room temperature, primary antibodies (see antibody table) were incubated at 4°C overnight, and horseradish peroxidase (HRP) conjugated secondary antibodies for 1 hour at room temperature. Membranes were exposed used SuperSignal^TM^ West Pico PLUS chemiluminescent substrate (Thermo scientific, 34580).

#### Statistical methods

Specific number of biological replicates and statistical methods used are specified in each figure or figure legend. IUE quantifications, single or double fluorescently labeled cells were quantified for at least three consecutive sections in each brain and their results averaged. To measure cell distribution in the cortex we used RapID ^40^. Briefly, a grid containing equally sized bins, numbers of bins specified in each figure, was manually placed in the cortex. The quantification of fluorescently labeled cells was automatically determined by the software. All statistical analyses and plot generation were performed using Prism 9 (GraphPad).

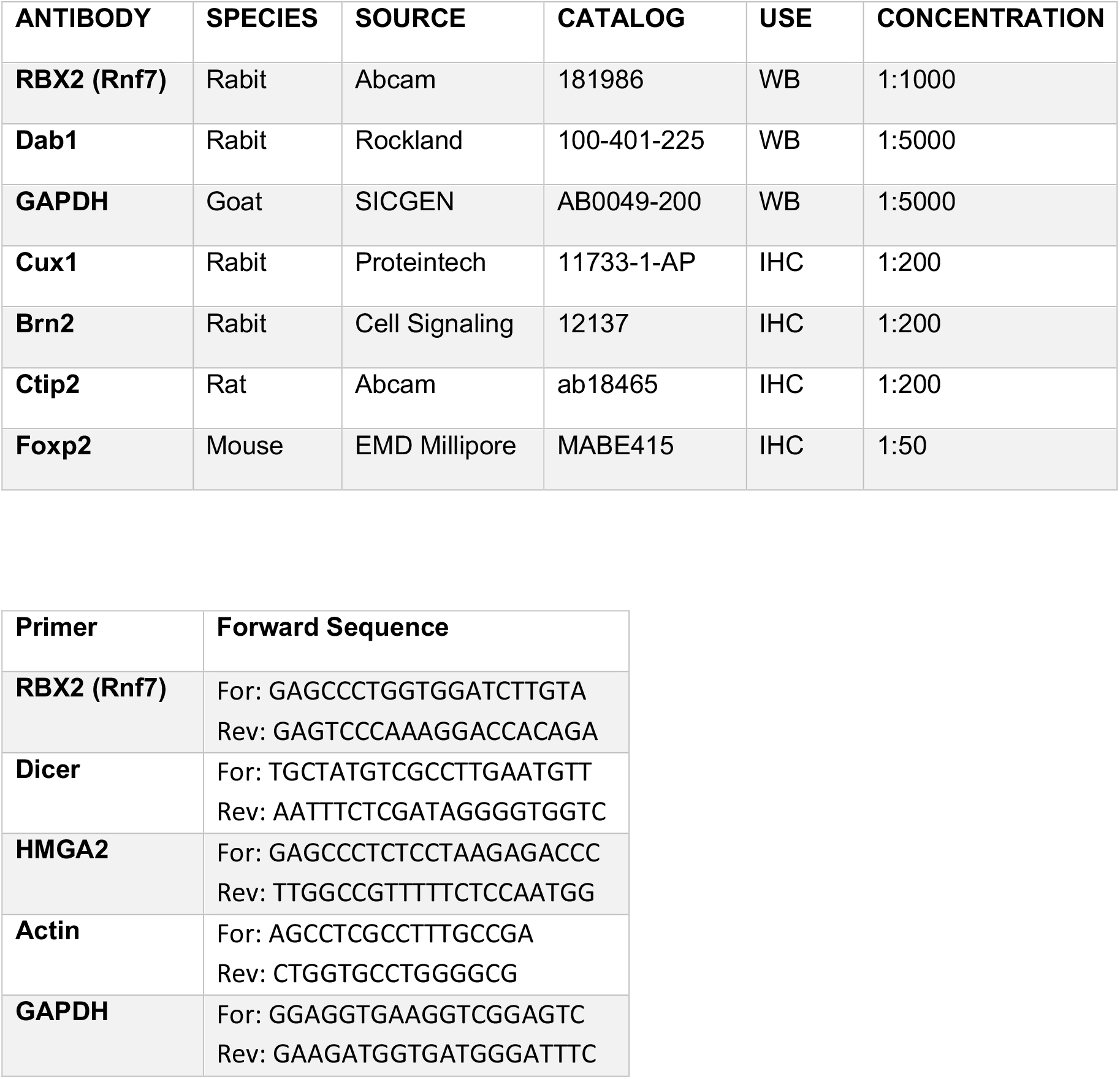

## Acknowledgements

We thank all members of the La Torre and Simó laboratories for their helpful insights. We also thank Drs. Tom Glaser and Nadean Brown for valuable comments and generosity with reagents. This study was supported by R01 NS43205 and R21 NS133564 to ALT and SS, and R01 NS109176 and R01 NS131375 to SS. We benefited from the use of the National Eye Institute Core Facility for histology sample processing and lentiviral production [supported by P30 EY012576]. The sequencing library preparations and the sequencing were carried out at the UC Davis Genome Center DNA Technologies and Expression Analysis Core, supported by NIH Shared Instrumentation Grant 1S10OD010786-01 and analyzed by the UC Davis Bioinformatics Core.

## Notes

### Competing Interest Statement

The authors have declared no competing interest.

