## Supplemental Figure 1 for "Uncoupling Neocortical Neuron Fate and Migration via a Let-7–RBX2 Axis"

A

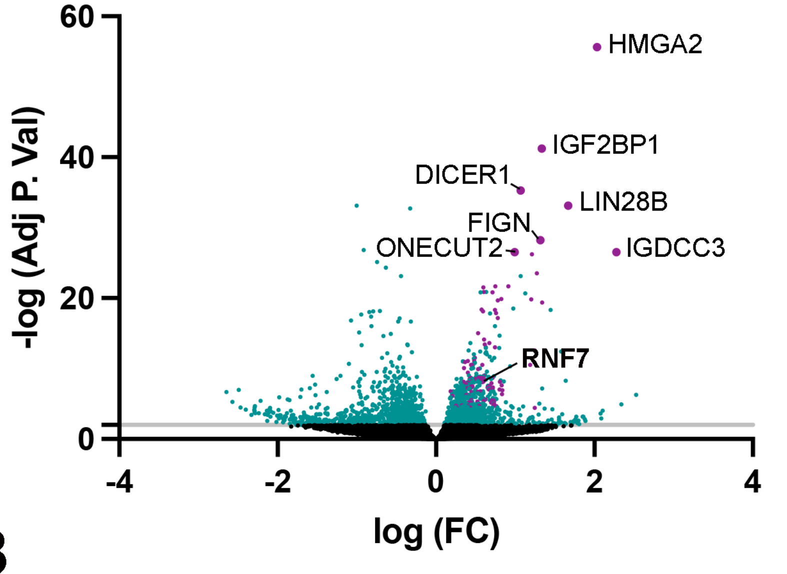


B

| Index | Name | P-value | Adjusted P-valu |
|---|---|---|---|
| 1 | G1/S Transition of Mitotic Cell Cycle (GO:0000082) | 4.40607E-06 | 0.0025 |
| 2 | Cell Cycle G1/S Phase Transition (GO:0044843) | 5.28138E-06 | 0.0025 |
| 3 | Negative Regulation of DNA-templated Transcription (GO:0045892) | 1.22823E-05 | 0.0038 |
| 4 | Negative Regulation of DNA-templated Transcription (GO:0045892) | 1.22823E-05 | 0.0038 |
| 5 | CRD-mediated mRNA Stabilization (GO:0070934) | 3.40961E-05 | 0.0080 |
| 6 | miRNA Processing (GO:0035196) | 5.98899E-05 | 0.0086 |
| 7 | Regulation of Gene Expression (GO:0010468) | 6.15118E-05 | 0.0086 |
| 8 | Regulation of Translation (GO:0006417) | 6.39692E-05 | 0.0086 |
| 9 | pre-miRNA Processing (GO:0031054) | 8.00738E-05 | 0.0094 |
| 10 | Mitotic Cell Cycle Phase Transition (GO:0044772) | 0.000113588 | 0.0119 |
| 89 | Regulation of Cell Migration (GO:0030334) | 0.015481662 | 0.1652 |
| 520 | Regulation of Neuron Migration (GO:2001222) | 0.188243931 | 0.3367 |
