## Supplementary figures and images for "Uncoupling Neocortical Neuron Fate and Migration via a Let-7–RBX2 Axis"

### Supplemental Figure 1

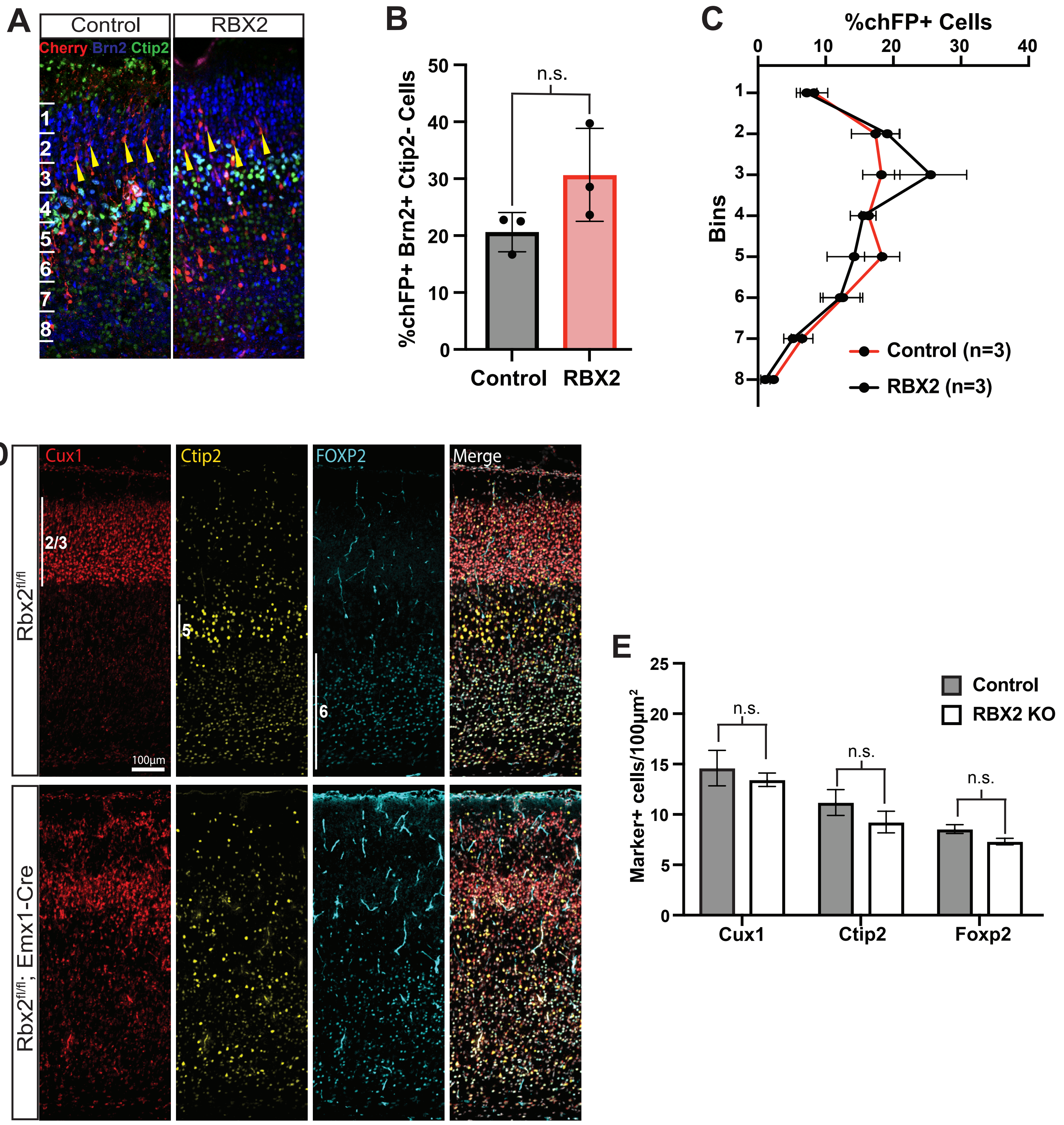

### Supplemental Figure 1

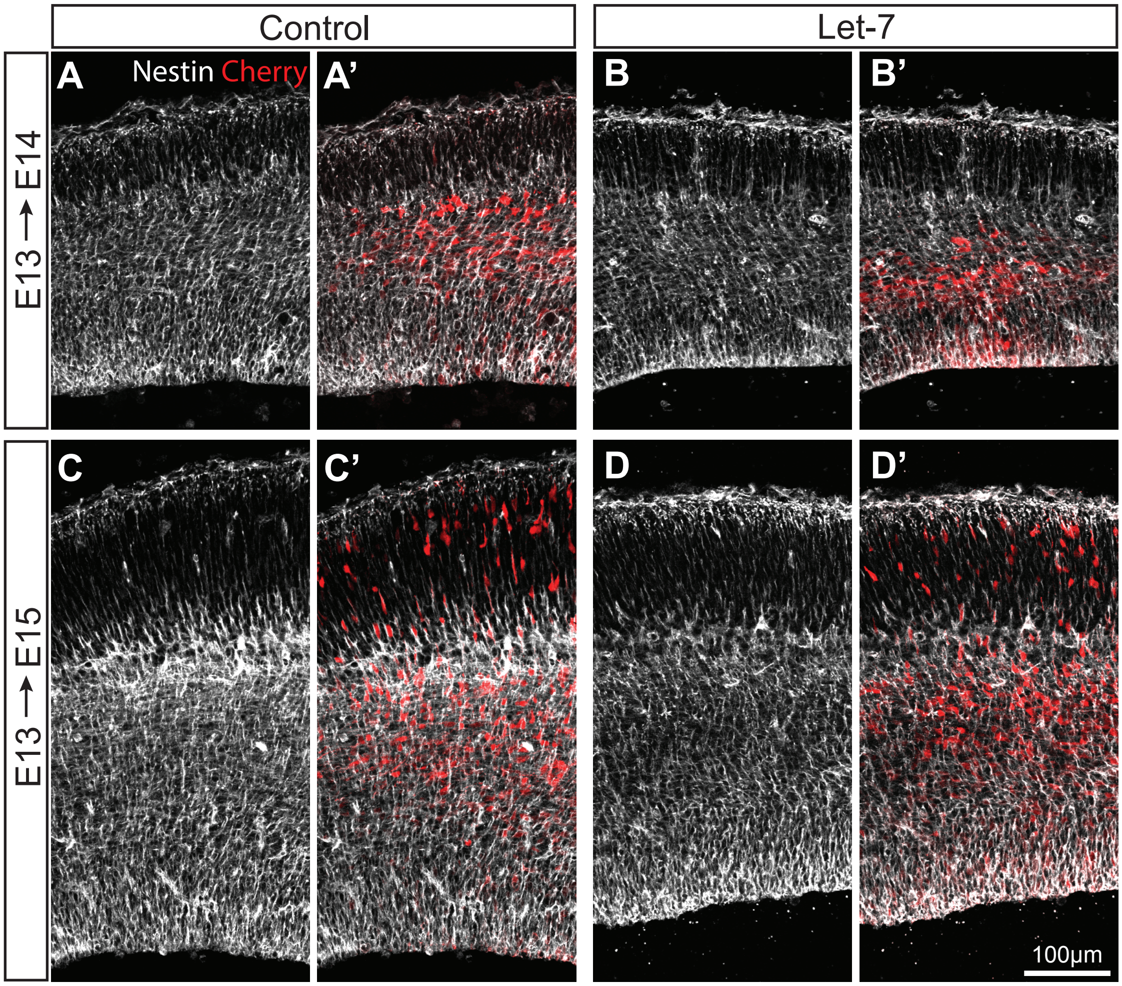
